# Revealing dynamic protein acetylation across subcellular compartments

**DOI:** 10.1101/472530

**Authors:** Josue Baeza, Alexis J. Lawton, Jing Fan, Michael J. Smallegan, Ian Lienert, Tejas Gandhi, Oliver M. Bernhardt, Lukas Reiter, John M. Denu

## Abstract

Protein acetylation is a widespread post-translational modification implicated in many cellular processes. Recent advances in mass spectrometry have enabled the cataloging of thousands of sites throughout the cell, however identifying regulatory acetylation marks have proven to be a daunting task. Knowledge of the kinetics and stoichiometry of site-specific acetylation are important factors to uncover function. Here, an improved method of quantifying acetylation stoichiometry was developed and validated, providing a detailed landscape of dynamic acetylation stoichiometry within cellular compartments. The dynamic nature of site-specific acetylation in response to serum stimulation was revealed. In two distinct human cell lines, growth factor stimulation led to site-specific, temporal acetylation changes, revealing diverse kinetic profiles that clustered into several groups. Overlap of dynamic acetylation sites among two different human cell lines suggested similar regulatory control points across major cellular pathways that include splicing, translation, and protein homeostasis. Rapid increases in acetylation on protein translational machinery suggest a positive regulatory role under pro-growth conditions. Lastly, higher median stoichiometry was observed in cellular compartments where active acetyltransferases are well-described.

## Introduction

Protein lysine acetylation is now acknowledged as a widespread modification, rivaling phosphorylation in scope^1^. Protein acetylation was first characterized on the N-terminal lysine residues of histone proteins that wrap DNA in the nucleus^2, 3^ and has since been described throughout the cell including cytoplasm^4^, mitochondria^5^, endoplasmic reticulum^6^, and peroxisomes^7^. In the nucleus, histone acetylation is associated with active gene expression, acting in part to open chromatin and allowing access for transcriptional machinery. Acetylation of cytoplasmic proteins affects diverse cellular processes which include cell migration, cytoskeleton dynamics, metabolism, and aging. Mitochondrial protein acetylation has been linked to metabolic regulation, oxidative stress, OXPHOS, and mitochondrial gene expression^8^.

Reversible acetylation is catalyzed by the enzymatic activity of lysine acetyltransferases (KATs) and deacetylases (KDACs), however other evidence also suggests that a significant proportion of acetyl-lysine sites result from nonenzymatic mechanisms^9–11^. We previously measured second-order rate constants for nonenzymatic acetylation *in vitro* and found that the chemical conditions of the mitochondrial matrix was sufficient to drive considerable nonenzymatic acetylation, and was generally consistent with lysine sites with low *in vivo* acetylation stoichiometry^12^. With the use of a general-base catalyst in the active site of KAT families that include GCN5 and MYST, these reactions are not affected by the protonation state of the ε-amino group, whereas the nonenzymatic reactions are directly dependent on the amount of unprotonated lysine^13, 14^.

Mass spectrometry has enabled the identification of over 20,000 acetylation sites in human cells^15^. This comprehensive catalog was facilitated by the development of antibody enrichment strategies for acetylated peptides^16, 17^. While immunoenrichment of acetylated peptides helps with identification of acetyl-lysine sites on low abundant proteins, immunoenrichment can introduce potential quantification bias through the additional experimental steps and antibody selectivity biases.

Unlike a plethora of examples from protein kinase signaling pathways, lysine acetylation has not been associated with analogous cascades, where one acetylation event of an acetyltransferase leads to acetylation of a second acetyltransferase to transmit a biological signal. Such cascades are used to amplify and rapidly propagate information down a signal transduction pathway. From available evidence, acetylation can modulate protein-protein and protein-DNA interactions, cellular localization, enzyme activity and stability^8^. For example, bromodomain-containing proteins recognize and bind acetyl-lysine residues for recruitment of larger multi-subunit complexes and permit efficient activation of gene transcription^18, 19^. In mitochondria, current evidence suggests that protein acetylation generally serves as an inhibitory modification of metabolic enzymes^8^. In this regard, acetylation appears to function as a rheostat to modulate the degree of a biochemical process. Given these examples of regulation, quantifying the level of stoichiometry is critical for understanding the biological effect of acetylation.

Protein phosphorylation cascades are well-known mechanisms by which cells respond in seconds to minutes to external stimuli. However, the time scales at which protein acetylation occur is poorly understood. Determining both stoichiometry and dynamic responses during cellular stimulation are key features to understand the role of protein acetylation at a site- and protein-specific level. Additionally, understanding how acetylation changes occur across subcellular compartments can provide insights into the cellular mechanisms controlling dynamic acetylation. As a part of this study, we provide an improved method using data-independent acquisition (DIA) to quantify acetylation stoichiometry at the proteome level, which was benchmarked using cellular proteomes with defined acetylation stoichiometry. We employed this method to understand how acetylation and proteome dynamics are concomitantly modulated in cells across subcellular compartments. Two human cell lines were synchronized by serum depletion/refeeding and monitored for changes in the proteome and site-specific acetylation, revealing rapid and dynamic changes in acetylation and protein expression profiles. Quantifying acetylation stoichiometry dynamics will be a critical tool for prioritizing the ever-increasing number of detected lysine acetylation sites for further investigation and towards a deeper understanding of this regulatory modification.

## Experimental Section

### Experimental Design and Statistical Rationale

#### Samples

For NCE optimization, MCF7 cultured cells were acquired using eight NCE parameters as described. For the acetylation stoichiometry calibration curve, a cell culture stock of HEK293 cells was used to prepare the light and heavy acetyl-modified samples which are combined into eleven mixed samples at varying amounts as described. Steady-state and dynamic stoichiometry analysis (MCF7 and HCT116 cells) were measured in biological triplicate.

#### Statistical tests

Various statistical tests including linear regression analysis and nonparametric analysis were performed throughout the study. Linear regression was performed on the stoichiometry calibration curve, while nonparametric tests were used to compare changes in the distribution of acetylation stoichiometry.

### Cell Culture conditions

MCF7 and HEK293 cells were grown using DMEM supplemented with 10% FBS. For global acetylation stoichiometry and single amino acid analysis, MCF7 and HEK293 cells were harvested at ∼80% confluency. Four hours prior to harvesting, cells were washed with PBS and replaced with fresh media. MCF7 and HEK293 cells were cultured for a total of 48 hours.

For the serum stimulation experiments, MCF7 and HCT116 cells were grown in DMEM supplemented with 10% FBS to ∼60% confluency. The cells were replenished with fresh media containing serum one hour prior to serum deprivation. The cells were then washed with PBS and replaced with fresh media without serum for 24 hours. After 24 hours, the 0 hour time point (no serum) was harvested and DMEM supplemented with serum was added to the other time points (15 min, 1, 2, or 4 hours).

All cells were harvested in 100 mM ammonium bicarbonate supplemented with acetyltransferase inhibitors (TSA, NAM, and sodium butyrate), protease inhibitors (PMSF, aprotinin, and leupeptin), and a phosphatase inhibitor cocktail (DOT Scientific).

### Sample preparation

We followed the procedure outlined in Lindahl *et al*, with the following exceptions described below^60^.

#### Protein chemical acetylation and digestion

Equal amount of protein (200 µg) was resuspended into 25-30 µL of urea buffer (8 M urea (deionized), 500 mM ammonium bicarbonate pH = 8.0, 5 mM DTT). Incubation steps throughout the sample preparation are carried out using the Eppendorf ThermoMixer® C. Sample was incubated at 60 °C for 20 minutes while shaking at 1500 RPM. Cysteine alkylation was carried out with 50 mM iodoacetamide and incubating for 20 minutes. Chemical acetylation of unmodified lysine residues was performed as previously described^12, 20, 34^. Briefly, ∼20 µmol of the “light” ^12^C-acetic anhydride (Sigma) or “heavy” D_6_-acetic anhydride (Cambridge Isotope Laboratories) was added to each sample and incubated at 60 °C for 20 minutes at 1500 RPM. The pH of each sample was raised to ∼8 using ammonium hydroxide and visually checked with litmus paper. Two rounds of chemical acetylation were performed for each sample to ensure near-complete lysine acetylation. To hydrolyze any O-acetyl esters formed during the chemical acetylation, the pH of the sample raised to ∼8.5 and each sample was incubated at 60 °C for 20 minutes at 1500 RPM. For protein digestion, the urea concentration of each sample was diluted to ∼2 M by adding 100 mM ammonium bicarbonate pH = 8.0 followed by addition of trypsin (Promega) at a final ratio of 1:100. The sample was digested at 37 °C for 4 hours while shaking at 500 RPM. If a second digestion using GluC (Promega) occurred, the urea concentration was further diluted to ∼1 M using 100 mM ammonium bicarbonate pH = 8.0 and digested with GluC (1:100) at 37 °C overnight while shaking at 500 RPM. Each sample was acidified by the addition of 15 µL of acetic acid.

#### Digesting protein sample to single amino acids

For complete digestion of proteins, which converts all unmodified lysine residues to free lysine, and all N-ε-acetylated lysine residue to acetyl-lysine, 20 μg of sample was diluted into 50 μL of digestion buffer (50 mM ammonium bicarbonate, pH 7.5, 5 mM DTT, in LC-MS grade water). A sample with 50 μL digestion buffer without protein was also included as a procedural blank. The samples were digested to single amino acids by treatment with three enzymes sequentially: First, samples were treated with 0.4 μg Pronase and incubated for 24 hr at 37°C. Then the Pronase activity was stopped by heating to 95°C for 5 min. After cooling down to ambient temperature, samples were then treated with 0.8 μg aminopeptidase and incubated at 37°C for 18 hr. Aminopeptidase activity was again stopped by heating samples to 95°C for 5 min and cooling down. Finally, samples were digested with 0.4 μg prolidase and incubated at 37°C for 3 hr. To extract the resulting single amino acids, 200 μL LC-MS grade acetonitrile (ACN) was added to each sample. The mixture was vortexed for 5 sec, spun at maximal speed for 5 min, and the supernatant was saved for analysis by LCMS.

#### Offline High pH Reverse Phase (HPRP) Prefractionation

Chemically acetylated peptides were resuspended into ∼2mL of HPRP buffer A (100 mM Ammonium Formate pH = 10) and injected onto a pre-equilibrated Phenomenex Gemini® NX-C18 column (5µm, 110Å, 150 × 2.0mm) with 2% buffer B (10% Buffer A, 90% acetonitrile). Peptides were separated with a Shimadzu LC-20AT HPLC system using a 2% – 40% buffer B linear gradient over 30 minutes at 0.6 mL/min flow rate, collecting 24 fractions throughout the length of the gradient. Fractions were dried down using a speedvac and pooled by concatenation into 6 final fractions as described previously ^61^.

#### Immunoblot Analysis

Samples were denatured in SDS loading dye and 5 min boil. 20 μg of whole cell lysate were loaded on a Bolt 4-12% Bis-Tris Plus gel (Invitrogen), followed by transfer to PVDF. Membrane was blocked with 1% milk before being blotted with either anti-Phospho-Akt (Ser473) (CST #4060) or anti-Akt (pan) (CST #2920) 1:2,000 and imaged. Bands were quantified using Image Studio Lite (LI-COR Biosciences).

### Mass spectrometry

#### Liquid chromatography

Peptides were separated with a Dionex Ultimate 3000 RSLCnano HPLC using a Waters Atlantis dC18 (100 µm × 150 mm, 3µm) C18 column. The mobile phase consisted of 0.1% formic acid (A) and acetonitrile with 0.1% formic acid (B). Peptides were eluted with a linear gradient of 2 – 35% B at a flow rate of 700 nL/min over 90 minutes. Peptides were injected by nanoelectrospray ionization (Nanospray Flex™) into the Thermo Fisher Q Exactive™ Hybrid Quadrupole-Orbitrap™ Mass spectrometer.

#### Data-dependent acquisition mass spectrometry

For data-dependent acquisition (DDA), the MS survey scan was performed in positive ion mode with a resolution of 70,000, AGC of 3e6, maximum fill time of 100 ms, and scan range of 400 to 1200 m/z in profile mode. Data dependent MS/MS was performed in profile mode with a resolution of 35,000, AGC of 1e6, maximum fill time of 200 ms, non-overlapping isolation window of 2.0 *m/z*, normalized collision energy of 25, dynamic exclusion was set for 30 seconds, and a loop count of 20.

#### Data-independent acquisition mass spectrometry

For data-independent acquisition (DIA), the MS survey scan was performed in profile mode with a resolution of 70,000, AGC of 1e6, maximum fill time of 100 ms in the scan range between 400 and 1000 *m/z*. The survey scan was followed 30 DIA scans in profile mode with a resolution of 35,000, AGC 1e6, non-overlapping 20 *m/z* window, and NCE of 25 or 30. For both DDA and DIA methods, the source voltage was set at 2000 V, capillary temperature at 250 °C, and S-lens RF = 50.

#### LCMS analysis of single amino acids

The abundances of free lysine, acetyl-lysine, and other amino acids from completely digested protein samples were analyzed using a Thermo Fisher Q Exactive™ Hybrid Quadrupole-Orbitrap™ Mass spectrometer coupled to a Vanquish UHPLC system (Thermo). Samples are separated using a 5 μm polymer 150 2.1 mm SeQuant® ZIC®-pHILIC column, with the following gradient of solvent A (ACN) and solvent B (10 mM ammonium acetate in water, pH 5.5) at a flow rate of 0.3 mL/min: 0-2min, 10% solvent B; 2-14min, linearly increase solvent B to 90%; 14-17min, isocratic 90% solvent B; 17-20min, equilibration with 10% solvent B. Samples are introduced to the mass spectrometer by heated electrospray ionization using a HESI II source. Settings for the ion source are: 10 aux gas flow rate, 35 sheath gas flow rate, 1 sweep gas flow rate, 3.5 kV spray voltage, 320°C capillary temperature, and 300°C heater temperature. Analysis is performed under positive ionization mode, with scan range of 88–500 m/z, resolution of 70 K, maximum injection time of 40 ms, and AGC of 1E6.

To quantify absolute levels of lysine and acetyl-lysine, an external calibration curve was run in the same sequence with the experimental samples. Lysine standard ranges between 10 to 200 μM, and acetyl-lysine standard ranges between 0.5 to 10 μM. Signal from procedural blank was subtracted from samples.

### Data Processing

#### Generating ^12^C-AcK and D_3_-AcK Spectral Library

The spectral library consists of a catalog of high-quality MS/MS fragmentation spectra resulting from data-dependent acquisition (DDA) MS runs. For the MCF7 and HCT116 stoichiometry, we performed DDA runs on three MCF7 lysate samples which were chemically acetylated with ^12^C-acetic anhydride, digested with trypsin and GluC, followed by HPRP prefractionation (see above). Prior to MS analysis, iRT peptides (Biognosys) were spiked into each sample following manufacturer’s guidelines. Database search was performed using MaxQuant with Andromeda as the peptide search engine version 1.6.1.0 using lysine acetylation and methionine oxidation as variable modifications and cysteine carbamidomethylation as a fixed modification. Enzymes for digestion were set to trypsin, which cleaves after lysines and arginines, and GluC, which cleaves after glutamate in ammonium bicarbonate buffers^62^. We increased the maximum missed cleavages to 4, because our labeling scheme, which modifies all unmodified lysines, prevents cleavage from trypsin. PSM and Protein FDR were both set to 1% calculated by the target-decoy approach, per default settings. Decoy entries were created in MaxQuant by reversing the original protein sequences. Main search peptide mass tolerance was set to 4.5 ppm, per default settings. The DDA runs were searched against the Swiss-Prot reviewed sequence database downloaded from UniProtKB on 12/12/2017 (20244 entries). The MaxQuant search results were imported into Spectronaut to build the 12C-AcK library. The 12C-AcK spectral library was then exported as a spreadsheet, specifically Biognosys’ library format (.kit), from Spectronaut and imported into a custom spectral library modifier, which completes the spectral library for all combinations of light and heavy acetylated peptides. With this *in silico* approach to inflate the 12C-AcK library, every acetylated peptide precursor will be represented by 2^n^ versions differing in the number and position of heavy/light acetylated lysine, where n is the number of acetylation sites in the peptide. The spectral library was completed with the corresponding precursor m/z values and fragment m/z values. The most intense fragment ions selected from the initial MS2 spectrum were cloned to the other peptide precursor versions. All peptide precursor versions will have identical retention time and hence iRT was also cloned. MaxQuant result files and inflated library can be accessed through ProteomExchange via the MassIVE repository (ProteomExchange: PXD014453; MassIVE: MSV000084029).

#### DIA MS data analysis

Data from DIA-MS was analyzed using Spectronaut 10. Thermo raw files were converted to HTRMS files with the Spectronaut Raw to HTRMS converter using the default settings and input into Spectronaut. The Spectronaut default settings for quantitation were used with slight modification: Identification-Qvalue score was set to 0.1 and Workflow-Unify peptide peaks were selected. This will cause Spectronaut to use the same integration boundaries for all light/heavy versions of one acetylated peptide within one LC-MS run. This change in the workflow will instruct Spectronaut to select for a given acetylated peptide precursor the best signal (by q-value) of the 2^n^ versions in the spectral library (see above). With this workflow, Spectronaut will then transfer the integration boundaries of the best scoring peptide precursor to the other peptide precursors. Because all of the 2^n^ peptide precursor versions only differ by the number of heavy instead of light acetylated lysine the retention time is expected to be identical. The spectral libraries which were completed as described above for all the light/heavy peptide precursor versions were used with this workflow. A Spectronaut output file containing all the fragment ion peak areas along with the corresponding peptide and protein identification was exported and used to compute the lysine site stoichiometry. A list containing all the data categories used for downstream stoichiometry analysis is found in the supplemental information.

#### Stoichiometry data processing

Data processing was performed in R v3.5.0 (http://www.r-project.org/) using an in-house made R script, which is available in the supplementary information. The stoichiometry preprocessing pipeline consists of two major steps: quantifying fragment ion stoichiometry and natural abundance isotopic correction, as described below. R-scripts utilized to analyze the results presented here can be accessed through GitHub (DOI:10.5281/zenodo.3360892).

#### Quantifying site-specific stoichiometry

DIA MS measures multiple peptide fragment ion abundances so this approach allows for quantitation of multiple lysines within a peptide. Acetylation stoichiometry of unique lysine sites are quantified by matching light and heavy fragment ion pairs and using the equation:

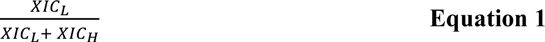

where XIC_L_ is the peak area of the light fragment ion and XIC_H_ is the peak area of the heavy fragment ion.

#### Isotopic purity correction

The mass shift of the light and heavy AcK peptides is 3 Da. This causes the M+0 peak of the heavy AcK peptide to overlap with the M+3 peak of the light AcK peptide. Therefore, we are correcting for the isotopic distribution overlap between the peptide pairs. This is done using an in-house R script as well as the R package, BRAIN v1.16.0 (Baffling Recursive Algorithm for Isotopic DistributioN calculations), available from Bioconductor, the open-source, software project (http://www.Bioconductor.org/)^63^. To correct for natural abundance of ^13^C isotope, the M+0 and M+I, where I represents the isotopic mass shift +1 or +3, were used to calculate the correction coefficient.

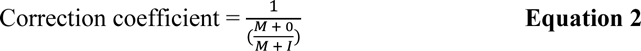

The correction coefficient is used to calculate the correction value:

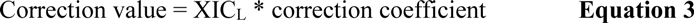

where the XIC_L_ is the peak area of the light fragment ion. Finally, the corrected heavy peak area (^Corr^XIC_H_) is calculated:

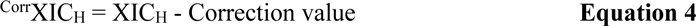

where the XIC_H_ is the peak area of the heavy fragment ion. The corrected stoichiometry is quantified using equation 1, substituting with ^Corr^XIC_H_.

#### MSstats

Protein abundance summarization was performed using MSstats v3.12.0 with the output of Spectronaut as the input. The function “SpectronauttoMSstatsFormat” was used with the following arguments: intensity set to “PeakArea”, filter_w_Qvalue set to TRUE, qvalue_cutoff set to 0.01, useUniquePeptide set to TRUE, fewMeasurements set to “remove”, removeProtein_with1Feature set to FALSE, and summaryforMultipleRows set to “max”. The dataProcesses function was then performed using the default arguments.

### NCE Optimization

To quantify site-specific acetylation stoichiometry from peptides containing multiple lysines, the fragmentation spectra of precursor ions must contain a high b- and y-ion coverage. To this end, we compared and optimized the number of peptide spectral matches (PSMs) as well as b- and y-ion coverage of MCF7 peptides (chemically acetylated with ^12^C-acetic anhydride followed by trypsin and GluC digestion) with a Q-Exactive MS using varying NCE settings (15, 20, 25, 30, 35, 40, 45, 50). For all NCE conditions, precursors between 400 – 1200 *m/z* were selected for fragmentation. MS1 resolution was set to 70,000, 3e6 target AGC, and 100 ms max IT in profile mode. MS2 resolution was set to 35,000, 1e6 target AGC, 200 ms max IT in profile mode with 15-sec dynamic exclusion. Database search was performed using MaxQuant version 1.5.4.1 followed by data analysis in R.

### Stoichiometry curve

We determined the accuracy and precision of the stoichiometry method by generating an 11-point stoichiometry curve using a complex sample. For this, we used a HEK293 lysate that was grown using standard culture conditions and harvested by centrifugation. The packed cell volume was resuspended using urea buffer (6-8M urea, 100mM ammonium bicarbonate pH = 8.0) and lysed by sonication. Protein concentration was measured using Bradford reagent (Bio-Rad).

To quantify stoichiometry ranging between 1-99%, we varied the amount of starting material to be chemically acetylated with ^12^C-acetic anhydride or D_6_-acetic anhydride using a total of 200 µg of protein for each stoichiometry point. For example, to measure a sample as 10% acetylated, we labeled 20 µg of HEK293 lysate with ^12^C-acetic anhydride and 180 µg of HEK293 lysate with D_6_-acetic anhydride. The starting protein amounts were varied to generate stoichiometries of: 1, 5, 10, 20, 40, 50, 60, 80, 90, 95, and 99% acetylation. Upon chemical acetylation, the sample was pooled together, digested using trypsin and we performed an offline HPRP prefractionation as outlined above.

### Bioinformatics

#### Subcellular localization assignment

To assign protein subcellular localization, we used the MitoCarta^64, 65^ and Uniprot (http://www.uniprot.org/) databases. For the “Mitochondrial” assignment of proteins, we used the Mitocarta database. Additionally, we used “Subcellular location” or “GO – Cellular component” from the Uniprot database to assign “Mitochondrial”, “Nuclear”, and “Cytoplasmic” pools. Other subcellular locations, such as endoplasmic reticulum, Golgi apparatus, cell membrane, etc., were assigned to the “Nuclear” fraction due to the likelihood that these cellular compartments, during differential centrifugation, would sediment in the “Nuclear” spin, which occurs at 1000 xg.

#### Quantitative Site set functional Score Analysis (QSSA)

The intersection of the KEGG pathway map^66^ and proteins in the spectral library detected with < 1% FDR was used for the gene set background. Acetylation coverage for each (p) pathway was calculated as the number of acetyl sites identified (*n_ack_*) over the total number of lysines in the pathway (*n_k_*), counted using protein sequences from Uniprot. The extent of acetylation was taken into account by summing the acetylation stoichiometry (s) across all conditions and all sites in each pathway. To allow for combining acetylation coverage and stoichiometry, the standard score of each quantity was taken. The overall pathway score was then calculated as the sum of the individual z-scores.

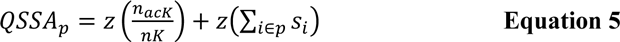

#### Functional analysis

Functional annotation of enriched gene ontology (GO) terms was assessed using DAVID v6.8^45, 46^. For enrichment analysis, the background was set to all the proteins identified in the DDA spectral library totaling 2400 unique protein IDs. The list of proteins with significantly changing acetylation sites in both serum-stimulated MCF7 and HCT116 cells were analyzed using DAVID for the following GO terms: GOTERM_BP_DIRECT, GOTERM_CC_DIRECT, and GOTERM_MF_DIRECT. GO term fold-enrichment was plotted as a bar graph with the corresponding p-value in overlaying, white text. Terms were grouped according to DAVID Functional Annotation Clustering.

#### STRING Network analysis

The underlying interaction network was downloaded from the STRING database (version 11.0)^47^. The thickness of edges in the STRING network display interaction confidence. Clusters of interactions were determined using k-means clustering with a set number of four clusters.

## Results

### DIA acetylation stoichiometry method optimization

We previously reported a method to determine lysine acetylation stoichiometry across an entire proteome^20^. This method employed an isotopic chemical acetylation approach to label all unmodified lysine residues within a sample and, upon proteolytic digestion coupled to LC-MS/MS, has been utilized to quantify proteome-wide acetylation stoichiometry in various biological conditions^20–23^ as well as in *in vitro*, nonenzymatic acetylation kinetics^12^. This method, however, had some limitations that we sought to address before utilizing this method to investigate acetylation dynamics. For example, 1) stoichiometry was measured using precursor-based quantification of the light and heavy peptide pairs. Therefore, signal interference in either the light or heavy channel could distort the stoichiometry quantification. 2) stoichiometry was quantified at the peptide level. Peptides can contain multiple lysine residues, so it was not possible to determine the contribution of individual lysines towards the observed stoichiometry. 3) software to analyze stoichiometry data is not widely available.

Here, we utilized an improved method to quantify acetylation stoichiometry in human cells by combining peptide prefractionation, data-independent acquisition (DIA) mass spectrometry, a novel spectral library generation procedure (**Figure 1A**). This method is applicable to both cell culture and tissue models because this workflow does not require metabolic labeling. A protein sample, extracted from either cell culture or tissue, is chemically acetylated using isotopic D_6_-acetic anhydride and digested with trypsin and GluC. The sequential digestion of the acetyl-proteome generates shorter peptides for MS analysis. Peptides are then pre-fractionated offline using high pH reversed-phase (HPRP) chromatography, analyzed using nano-LC-MS/MS in DIA mode and analyzed using Spectronaut (Biognosys)^24–27^ (**Figure 1A-right side, Figure S1**). A novel aspect of the method includes a project-specific spectral library that is generated from a D_0_-acetic anhydride (light isotope) modified sample. Acetyl peptides are pre-fractionated using HPRP and acquired in data-dependent acquisition (DDA) mode. The spectral library, which includes light acetyl peptides (D_0_-AcK), is imported into Spectronaut. To account for both light and heavy acetyl-lysine fragment ions in the spectral library, a novel, standalone software was developed to be used with Spectronaut, which generates the heavy labeled (D_3_-AcK) fragment ions *in silico* from the light spectral library (**Figure 1A-left side**). This process ensures that for a given peptide all light and heavy fragment ions are represented in the spectral library. This is a critical step for deeper coverage of the proteome, and supports a more rigorous analysis that requires both isotopic pairs to be quantified before stoichiometry is calculated. Combining offline pre-fractionation and DIA has several advantages that addresses unique limitations from the original study, as has been previously discussed^23, 24, 27–30^. HPRP prefractionation reduces interferences caused by coeluting peptides and has the added benefit of also increasing the depth of the acetylome coverage^31^. DIA analysis uses all quantified light and heavy fragment ions in MS2 which yields more accurate quantification than using MS1 signal. Additionally, stoichiometry can be localized to a specific lysine site even when multiple lysines are present on a peptide (**Figure 1B**).

**Figure 1:**
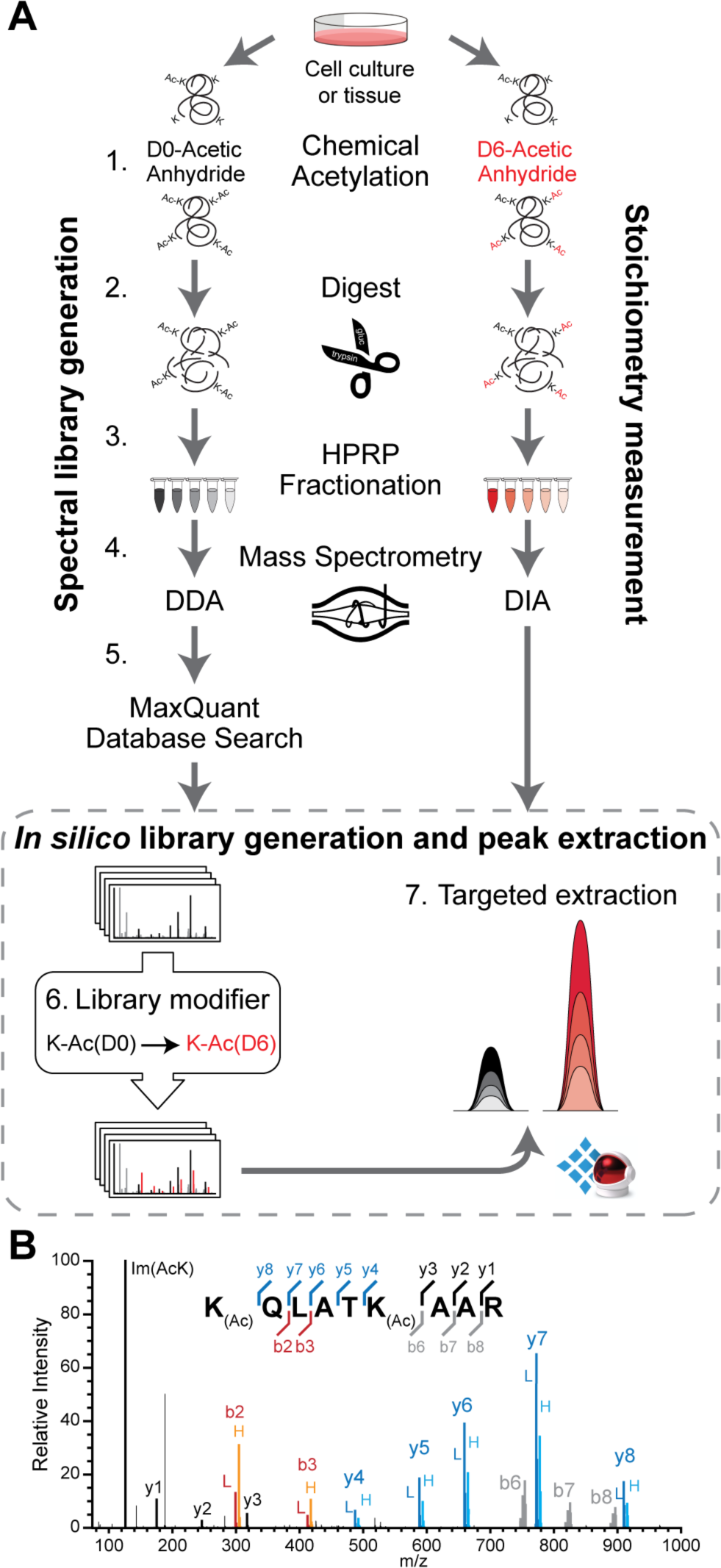
DIA acetylation stoichiometry workflow(A) Overall workflow for quantifying acetylation stoichiometry. A spectral library containing all possible light and heavy fragment ions is generated by 1) chemical acetylation of a sample using D_0_-acetic anhydride, 2) digestion using trypsin and GluC, 3) high pH reverse phase (HPRP) fractionation, 4) DDA mass spectrometry followed by a MaxQuant database search. The spectral library is imported into Spectronaut and using an external standalone software tool, the spectral library is 5) modified to contain all the heavy acetyl-lysine fragments. To quantify stoichiometry, the sample is 1) chemically acetylated using D6-acetic anhydride, 2) digested, 3) pre-fractionated 4) analyzed using DIA mass spectrometry and 6) analyzed in Spectronaut for targeted extraction using the custom light and heavy acetyl-lysine spectral library. (B) Diagram illustrating MS2 spectra for the Histone H3 peptide containing lysine K18Ac and K23Ac. The fragments b2-b3 are specific for K18 and y4-y8 are specific for K23. The fragments b6-b8 are ambiguous as they contain K18Ac and K23Ac, while the fragments y1-y3 contain no acetyl lysine information.

To assess the accuracy and precision of the improved workflow, a proteome-wide acetylation stoichiometry calibration curve was generated. For this analysis, a proteome sample of HEK293 cells was chemically acetylated with either light (^12^C-) or heavy (D_6_-) acetic anhydride, which were then combined at varying ratios and subjected to the DIA workflow (**Figure 2A**). A cell-based proteome was used for method validation over standard peptides, which had several advantages: *1*.) The use of standard peptides at this scale would be cost prohibitive. *2*.) The combined acetyl-proteomes results in comprehensive acetylation stoichiometry, mimicking experimental samples. *3*.) The level of acetylation can be modulated to encompass a wide range of stoichiometries (1% – 99%) providing limits of sensitivity and quantification. A caveat of using a cell-based workflow is that endogenous acetylation can potentially confound the results, which can lead to overestimation of some lysine sites. However, because the vast majority of lysine sites display negligible acetylation, this validation approach can be used to assess the accuracy and precision of the stoichiometry workflow.

**Figure 2:**
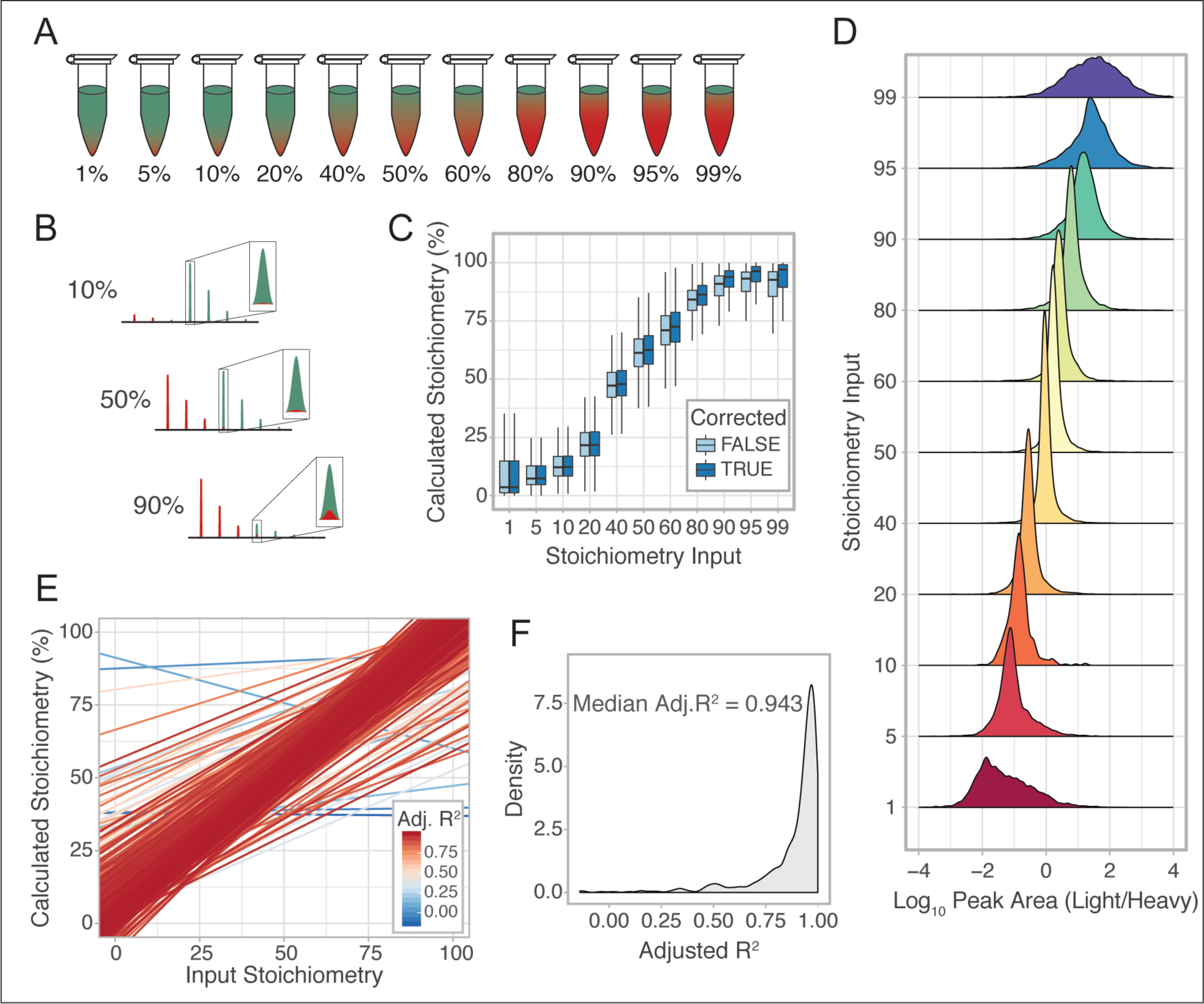
Benchmarking the DIA stoichiometry workflow (A) Diagram of standard curve samples generated by combining fully heavy labeled (green) peptides with fully light labeled (red) peptides at the specified ratios. (B) Correction of natural abundance isotopic envelope between light and heavy acetyl peptides. Red peaks represent light acetyl containing fragment ions and the green peaks represent heavy acetyl fragment ions. Isotopic overlap is much more pronounced at high stoichiometry. (C) Boxplot of the proteome-wide stoichiometry distribution before and after natural abundance isotope correction. (D) Density plot of the Log_10_-transformed (light/heavy) peak area for each input stoichiometry condition. (E) Linear regression analysis of all acetyl lysine sites (> 9 data points, n = 616) with the color of each line corresponding to the R^2^ value. (F) Distribution of Adjusted R^2^ values for the acetyl lysine linear regression analyses.

Due to the 3 Da mass shift between the light and heavy acetyl peptide fragment ions, high acetylation stoichiometry will lead to an underestimation of stoichiometry due to increased intensity of the M+3 natural abundance isotopic peak (**Figure 2B, Figure S2**). To account for this, a natural abundance correction was applied to all heavy acetyl lysine fragment ions. This correction was performed by subtracting the M+3 isotopic peak of the light acetyl lysine fragment ion from the M+0 isotopic peak of the heavy acetyl lysine fragment ion (**Figure 2B, Figure S2**). This global correction improved the precision of the stoichiometry quantification, especially in the higher stoichiometry values (**Figure 2C**). An alternative method to assess the precision of the quantification is to measure the ratio of the light and heavy fragment ions. Quantification of the light/heavy ratios corresponding to stoichiometry profiles between 20 and 80% displayed the highest precision (**Figure 2D**). This is due to the abundance values of the light and heavy fragment ions near 1:1 ratio. In contrast, stoichiometries at the extreme ends of the curve (< 5% and > 95%) displayed the lowest precision since quantitation in these conditions requires the measurement of fragment ions greater than 20-fold difference (**Figure 2D**). In order to calculate stoichiometry, both the heavy and light fragment ions must be observed, which can be challenging for the lowest stoichiometry if the light fragment ion is below the limit of detection. The heavy fragment ions or light fragment ions are summed independently before calculating the stoichiometry. This allows the higher intensity, higher confidence fragment ions to strongly influence the stoichiometry calculation, rather than biasing the calculation by equally weighting high and low abundance fragment ions. The stoichiometry curve analysis quantified stoichiometry for ∼1400 acetyl lysine sites. The number of acetyl sites quantified in at least nine conditions was 616. Linear regression analysis using acetyl lysine sites quantified in at least nine conditions shows high reproducibility of this method (**Figure 2E**) with a median R^2^ of 0.94, after correction for multiple regression analysis (**Figure 2F**). This global analysis with well-defined input stoichiometries highlights the quantitative nature of this method and is applicable to query acetylation stoichiometry of an entire proteome.

The improved stoichiometry workflow enables the quantitation of different acetyl-lysines from a single peptide, removing the ambiguity of site quantification. As an example, the histone H3 peptide (containing K18 & K23), K_Ac_QLATK_Ac_AAR, has fragment ions that are unique to each lysine site. K18 is quantified by the fragment ions b2-b3, while y4-y8 are specific for K23 (**Figure 1C**). Obtaining high quality and high coverage of b- and y-ions is essential for quantification of multiple lysines on the same peptide. Therefore, the normalized collision energy (NCE) was optimized for higher energy collisional dissociation (HCD) fragmentation ^32^. Peptide spectral matches (PSMs) were evaluated (**Figure 3A**) as well as the global b- and y-ion coverage (**Figure 3B**) across a wide range of NCEs (15-50 in 5-unit increments) using a chemically acetylated, trypsin and GluC digested proteome. A low number of PSMs with a c-terminal lysine are observed (blue bar). These peptide matches could arise from trypsin cleavage of unmodified lysines as well as proteins with a c-terminal lysine. However, comparing the frequency of lysine peptides to c-terminal glutamate (red) and arginine (green) peptides demonstrates that the chemical acetylation of the proteome progresses to near completion. To determine the global b- and y-fragment ion coverage, each fragment ion identified for a given PSM was counted and normalized to the peptide length. As y-ions increase with higher NCE, the proportion of b-ions begin to decline at a similar rate (**Figure 3B**). The NCE 25 was used to balance the frequency of b-ions, y-ions, as well as the number of PSMs (**Figure 3A**).

**Figure 3:**
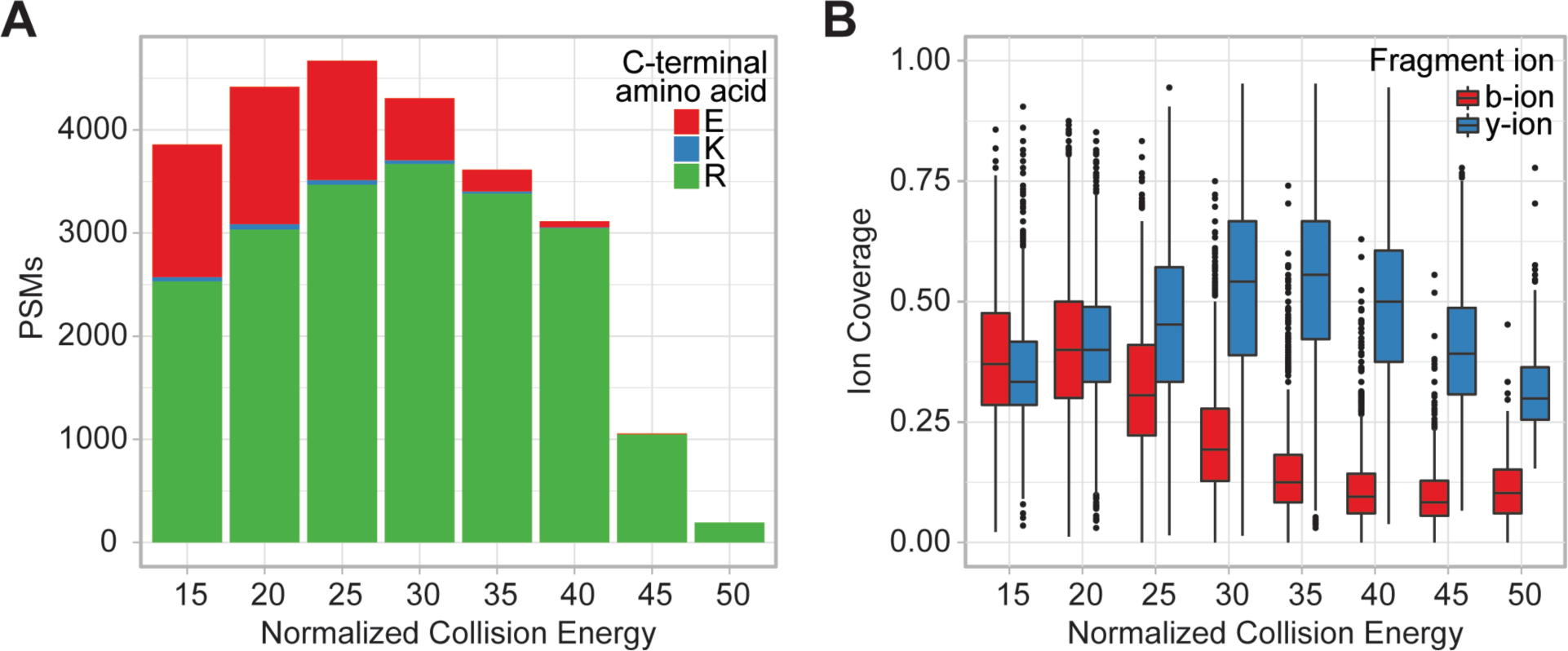
Optimization of instrument parameters for increased fragment ion coverage (A) Peptide spectral matches (PSMs) at varying normalized collision energy (NCE) for trypsin (c-terminal arginine and lysine) and GluC (c-terminal glutamate) digested peptides. The data consists of an MCF7 whole cell lysate labeled with 12C-acetic anhydride, digested with trypsin and GluC and analyzed in DDA mode using varying NCE settings. (B) Global fragment ion coverage for b- and y-ions of the data in A.

### Subcellular distribution of acetylation stoichiometry

There are few studies measuring (or estimating) acetylation stoichiometry in mammalian systems^21, 22, 33^. Thus, a comprehensive analysis of the acetylation stoichiometry distribution across the cell remains uncertain. To address this, we first utilized the current quantitative stoichiometry approach using breast cancer cell line MCF7 and quantified a wide range of stoichiometry (< 1% up to 99%) with high correlation between acetyl lysine fragment ions (red) and peptides (blue) between three biological replicates (**Figure 4A**). Quantifying acetylation stoichiometry in MCF7 cells shows the distribution of acetylation skewed towards low stoichiometry (**Figure 4A**). To determine if the distribution of stoichiometry varies across subcellular regions, we next grouped each protein into known subcellular localization based on Uniprot localization and compared the acetylation stoichiometry distribution between cytoplasmic and nuclear localizations. Using this grouping and non-parametric analysis, the nuclear fraction contains more acetylation sites with a higher stoichiometry compared to the cytoplasmic fractions (p = 0.00027) (**Figure 4B**). To validate these findings, an orthogonal approach to quantify subcellular acetylation levels was utilized. Subcellular fractionation was performed on MCF7 cells by differential centrifugation and acid extraction to enrich for histone, nuclear non-histone, mitochondrial, and cytosolic proteins. Each fraction was treated with a combination of proteases to completely digest proteins of each subcellular compartment to individual amino acids. The relative abundance of acetyl-lysine and unmodified lysine can be measured using mass spectrometry^34^ (**Figure 4C**). Acetyl-lysine was significantly more abundant on histone and nuclear proteins compared to the cytoplasm and mitochondrial fractions (**Figure 4D**) corroborating the peptide-level stoichiometry results (**Figure 4B**).

**Figure 4:**
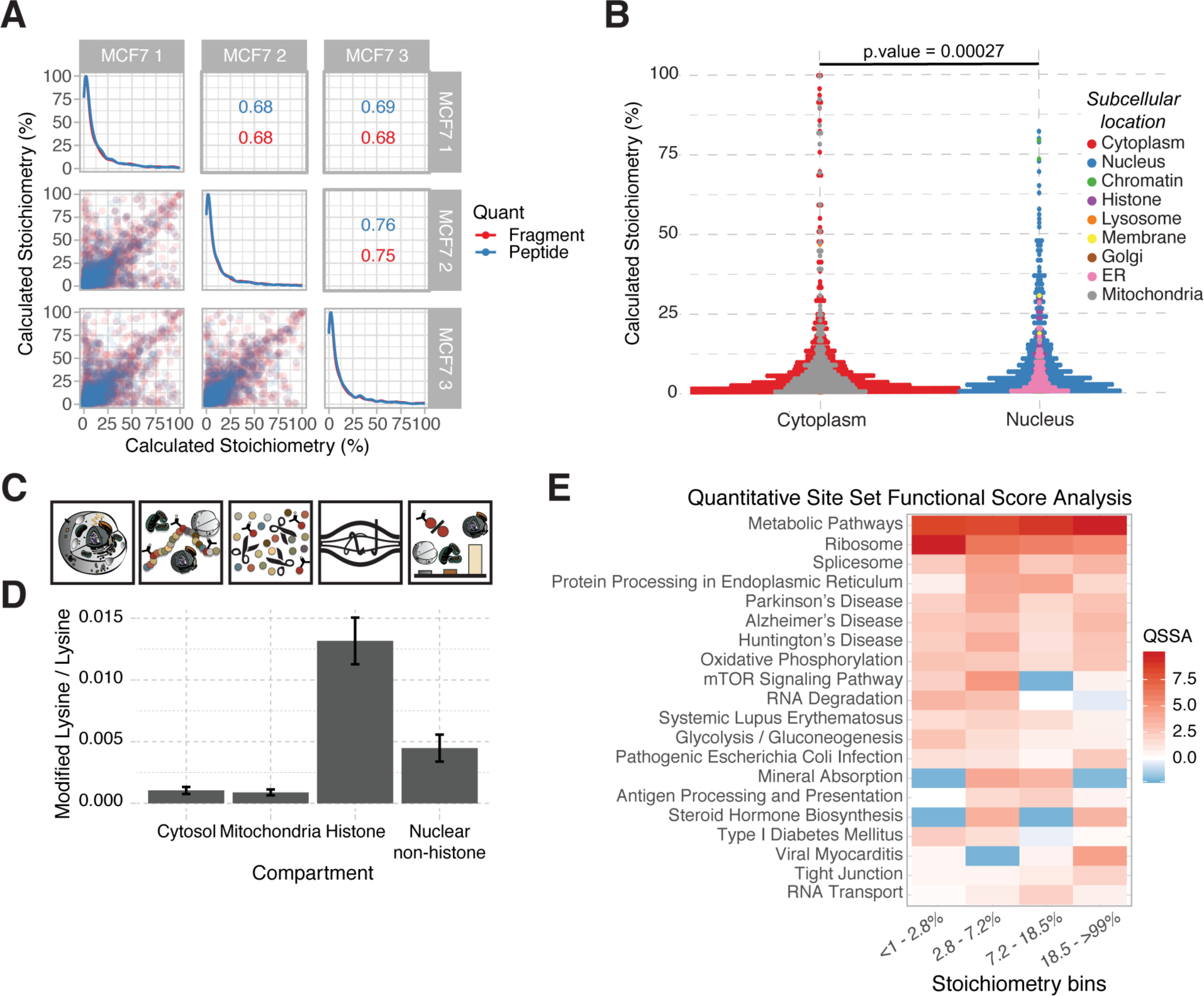
Nuclear-localized proteins have the highest lysine PTM abundances (A) Scatterplot matrix of acetylation stoichiometry across the three biological replicates of MCF7 cells. Spearman correlation is displayed for each pairwise comparison of fragment ions (red) acetyl-lysine site stoichiometry (blue). (B) QSSA heatmap showing enriched KEGG pathways. Acetylation stoichiometry was binned into quartiles and used as input for the QSSA. (C) Acetylation stoichiometry distribution across cellular compartments. Compartments were grouped in silico as cytoplasmic or nuclear. Statistical analysis was performed using the posthoc Kruskal-Nemenyi test. Colored circles represent individual subcellular location assignment based on Uniprot location and mitocarta database 64,65. (D) Workflow for quantifying single amino acid abundances of modified and unmodified lysines of enriched subcellular fractions. Each fraction is digested with a cocktail of proteases to generate single amino acids. Amino acids are analyzed by MS to quantify modified lysine / unmodified lysine as a measure of global subcellular stoichiometry. (E) Acetyl-lysine:lysine ratio across cytoplasmic, mitochondrial, histone, and non-histone proteins.

To identify biological processes enriched in acetylation, a biological pathway analysis tool recently developed termed quantitative site set functional score analysis (QSSA) was used to analyze the acetylation stoichiometry dataset^35^. This tool was developed for PTM pathway enrichment analysis taking into account the number of modified sites as well as the fold-change across conditions. QSSA was adapted for acetylation stoichiometry datasets. The stoichiometry data from MCF7 cells was divided into quartiles (for stoichiometry ranges, see Materials and Methods). Each quartile was used as input for the QSSA. To correct for any sample preparation or mass spectrometry analysis biases, QSSA enrichment scores were calculated against a background of proteins and acetyl-lysine sites identified in the spectral library. Gene Ontology processes that were enriched in this experiment include Metabolic Pathways, Ribosome, Spliceosome, and Protein Processing in Endoplasmic Reticulum (**Figure 4E**). Enrichment of metabolic pathways is a hallmark of acetylation studies^3, 35, 36^. Proteins that form part of the ribosome are N-ε-^17, 37^ and N-α-acetylated^38^. Interestingly, decreases in N-α-acetylation of ribosomal proteins correlate with a decrease in 80S ribosome assembly and cell growth^38^ demonstrating a functional link, however, it remains unknown whether N-ε-acetylation can regulate ribosomal function or what time scales are needed to observe changes in ribosomal acetylation.

### Acetylation and proteome dynamics

While many studies focus on acetylation changes over longer periods of time, days in cell culture and months in animal models, understanding the dynamics of acetylation over shorter time scales (minutes and hours) can give critical insight into the mechanisms and functionality of acetylation. With time-course information, the relative timing, direction, and magnitude of the acetylation changes can help to distinguish change as primary or secondary responses. Levels of acetylation are dictated through additive mechanisms that involve acetyltransferase activity, nonenzymatic reactions and increased acetyl-CoA levels, while removal mechanisms involve deacetylase activity and protein degradation^8, 36, 39^. Therefore, understanding the dynamics of not only protein acetylation, but also proteome dynamics will be important to understand the interplay between these two processes. The majority of studies quantifying acetylation dynamics utilize an antibody based workflow to enrich for the acetyl-peptides^16, 17^. Using an enrichment strategy, it is necessary to account for changes in protein abundance in order to accurately report changes in acetylation by analyzing a sample of the proteome which was not subjected to the immuno-enrichment procedure. The methods described here represent a label-free DIA workflow that does not require an enrichment step. Instead, all free lysine residues are chemically modified using acetic anhydride, a step which is analogous to the alkylation of cysteine residues with iodoacetamide. Therefore, precursor abundance data collected from the acetylation stoichiometry workflow can also be used to estimate protein abundance using label-free quantification techniques.

To initiate a robust stimulation of MCF7 cells, 24 hr serum-starved cells were activated with serum and harvested at 0, 1, 2, 4 hours (**Figure 5A**). Acetylation stoichiometry and protein abundance were determined as in Figure 1A. This design has the benefits of synchronizing cells upon serum starvation followed by robust changes in signaling pathways that occur upon serum replenishment^40–42^. To verify activation of major signaling pathways, we monitored the level of AKT S473 phosphorylation (**Figure 5B, C**) and S6 ribosomal protein S235/236 phosphorylation (**Figure S3**).

**Figure 5:**
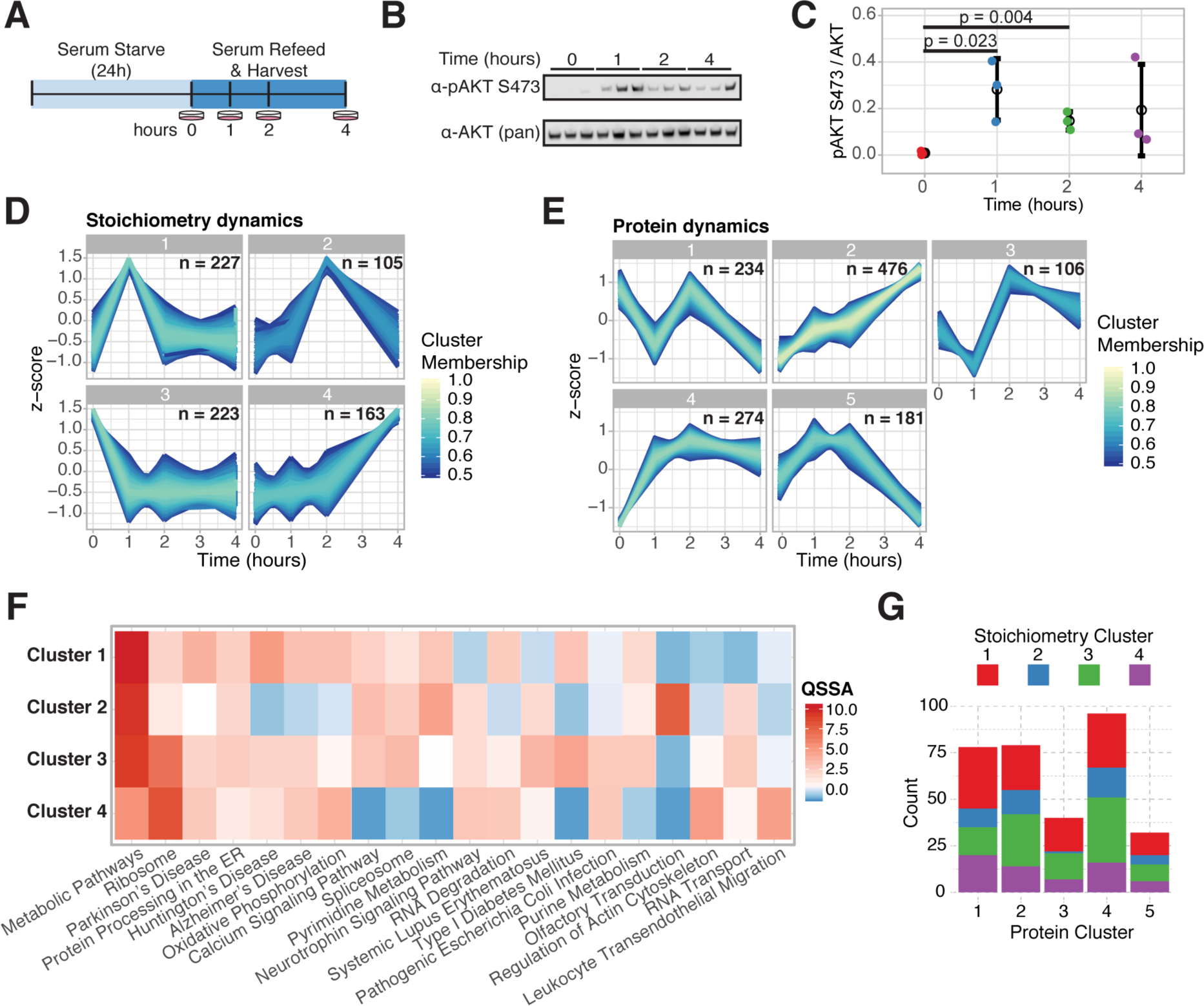
Acetylation and protein dynamics (A) Diagram of experimental approach to quantify acetylation stoichiometry dynamics. (B) Immunoblot of phosphorylated AKT S473 (top) and AKT (bottom). (C) Quantitation of the immunoblot. Statistical analysis was performed using a t-test. (D) Time-course clusters of acetylation stoichiometry dynamics by fuzzy c-means clustering. (E) Time-course clusters of protein abundance dynamics by fuzzy c-means clustering. (F) QSSA heatmap showing enriched KEGG pathways. Acetylation stoichiometry was separated by clusters and used as input for the QSSA. (G) Barplot quantifying the frequency of acetylation dynamic clusters within each protein dynamics cluster.

Acetylation stoichiometry was quantified using the described DIA-MS approach followed by a pattern recognition analysis using fuzzy c-means clustering^43^. Clustering analysis identified four unique clusters where site-level acetylation dynamics revealed distinct profiles (**Figure 5D**). Over two-thirds of the acetylation sites identified in this clustering analysis were found in clusters 1 and 3 which display rapid changes upon growth factor stimulation. These clusters correspond to acetylation levels that rapidly increased and returned to pretreatment baseline levels (Cluster 1) as well as trends where acetylation rapidly decreased and remained low (Cluster 3), respectively. Protein abundance was determined by label-free quantification using MSstats in conjunction with the chemically acetylated proteome sample^44^ followed by the clustering analysis using fuzzy c-means (**Figure 5E**)^43^. QSSA analysis of the serum-stimulated acetylation stoichiometry dataset identified biological processes that are enriched in each of the acetylation clusters **(Figure 5F)**. Acetylation clusters 1 and 2, which exhibited rapid increases in acetylation stoichiometry within the first two hours were highly enriched for Metabolic Pathways. Cluster 4, which contained acetylation sites that more slowly increased over time, was highly enriched for the Ribosome.

To investigate possible links between protein levels and acetylation dynamics, the trends of acetylation were compared within the protein clusters (**Figure 5G**). For example, there is strong overlap between acetylation stoichiometry cluster 3 within the protein clusters 2 and 4. This demonstrates that there is a subset of sites that have decreased acetylation stoichiometry with increasing protein abundance. This particular overlap of dynamics could be due to an increase in protein abundance without increased lysine acetylation, which would result in a trend showing a decrease in acetylation stoichiometry. Whereas, increases in lysine acetylation when protein abundance also increases must occur through active acetylation.

### Coordinated acetylation dynamics in diverse human cell lines

The results from MCF7 cells suggest protein acetylation is dynamically controlled in a model of growth factor stimulation, and that major metabolic and cellular pathways are targets of these rapid acetylation changes. To further understand acetylation dynamics and determine if the serum-stimulated changes are conserved across cell types, the serum starve-replete cell culture model was applied to a colon cancer cell line, HCT116 (**Figure 5A)**. Acetylation stoichiometry was quantified using the DIA-MS approach at 0 hours (no serum), 15 minutes, 1, 2, and 4 hours post-serum re-addition. The total number of quantified acetylation sites in the HCT116 experiment was 3818, which is comparable to the 4310 acetylation sites quantified in the MCF7 experiment.

Interestingly, there was a strong overlap of quantified acetylation sites across the two experiments, 3143 sites had quantifiable stoichiometry in both experiments resulting in a ∼63% overlap (**Figure 6A**). To identify and understand which acetylation sites are dynamic in both experiments, we identified which acetylation sites are significantly changing over time using one-way analysis of variance (ANOVA). There were 647 and 1037 acetylation sites in MCF7 and HCT116 cells, respectively, significantly changing over time (p-value ≤ 0.05) with 228 sites significantly changing in both experiments (**Figure 6B**). Observing dynamic acetylation on the same lysine site in diverse cell lines suggests functional conservation. Examples of overlapping acetylation sites are Mitochondrial fission 1 protein (FIS1) K89 and Sulfiredoxin (SRXN1) K116. Both of these acetylation sites experienced rapid increases in acetylation in response to serum stimulation, a 30-40% increase at FIS1 K89 and a 25-60% increase at SRXN1 K116, which was maintained at a high level throughout the time course.

**Figure 6:**
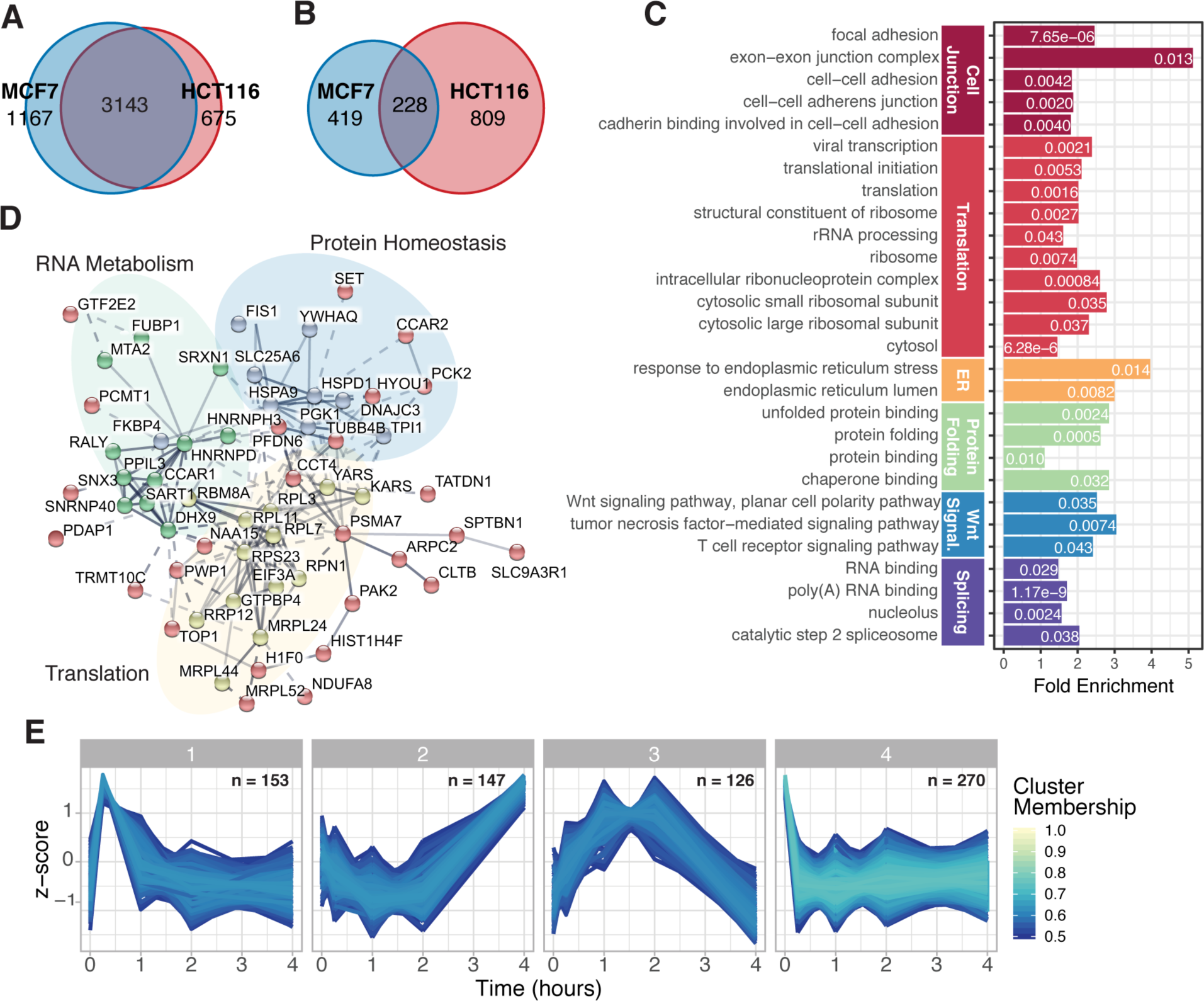
Coordinated acetylation dynamics in diverse cell lines, MCF7 and HCT116 (A) Venn diagram of quantified acetylation sites in serum-stimulated MCF7 and HCT116 cells. (B) Venn diagram of significantly changing acetylation sites (significance calculated using one-way ANOVA analyses) between MCF7 and HCT116 cells. (C) DAVID GO Term enrichment analysis of the proteins significantly changing in both MCF7 and HCT116 cells. Bar graphs represent fold enrichment with p-values in white text, and terms are grouped by similarity. (E) STRING network analysis of robustly (≥ 5%) and significantly changing sites in both experiments. Color of nodes represent cluster from k-means clustering in STRING and edges represent the confidence of the interaction. (F) Time-course clusters of acetylation stoichiometry dynamics by fuzzy c-means clustering.

In addition to identifying the proteins and sites significantly changing in both experiments, determining the pathways similarly regulated between cell lines was an important goal. Therefore, we used DAVID Bioinformatics Functional Annotation Tool to identify GO term enrichment in the overlapping dynamic acetylation sites^45, 46^. This enrichment was calculated against a background of identified proteins in our spectral library to correct for any sample biases. Grouping similar GO Terms together, we found that terms associated with cell junction, translation, protein folding, and splicing were enriched (**Figure 6C**). Furthermore, we took the most robustly changing sites in the overlap, ≥ 5% change up or down relative to the pre-stimulation time point and performed network analysis using STRING (**Figure 6D**)^47^. K-means clustering of the resulting network demonstrated three main clusters of proteins that correlated with the pathway enrichment analysis. The clusters generally encompassed splicing and RNA-binding proteins, translational machinery, and proteins involved in maintaining protein homeostasis.

Lastly, we used the same fuzzy c-means clustering to identify acetylation dynamic patterns in the serum-stimulated HCT116 cells (**Figure 6E**). Interestingly, similar acetylation trends in HCT116 cells were also observed in the MCF7 cells (**Figure 5D**). We identified several clusters with rapid changes in acetylation (clusters 1, 3, and 4) and one cluster with a delayed acetylation response (cluster 2). When we overlaid the pathway enrichment analysis onto the cluster analysis, we found that RNA metabolic processes, such as splicing and translation, had more sites that increased in acetylation over time (**Figure 5D – clusters 1 and 4, and Figure 6E – clusters 1, 2, and 3**). Cell-cell junction annotated proteins demonstrated more sites whose acetylation decreased over time with growth factor stimulation (**Figure 5D – cluster 3 and Figure 6E – cluster 4**).

## Discussion

### Integrated stoichiometry workflow

In this study, we have developed and utilized an improved DIA mass spectrometry method to quantify acetylation stoichiometry and investigate rapid acetylation dynamics in cells in response to serum stimulation. Understanding the speed and magnitude of protein acetylation changes on shorter timescales (minutes and hours) provide new insights into the cellular mechanisms of acetylation and help to prioritize specific sites for functional investigation. As discussed earlier, this improved method addresses limitations of our previously developed DDA method (20). Here, we integrated the acetylation stoichiometry workflow with targeted DIA analysis, which allows identification and quantification of light and heavy acetyl-lysine fragment ions, enabled by a novel spectral library that contains all light and heavy acetyl-lysine feature pairs. Using this method, we demonstrate accurate and reproducible analysis of dynamic protein acetylation in different cell lines and highlight similar activated pathways upon serum stimulation.

Recently, Choudhary and Colleagues developed an orthogonal method to quantify acetylation stoichiometry^30, 33, 48^, which incorporates a heavy SILAC labeled proteome subjected to partial chemical acetylation, combined with an experimental sample by serial dilution, trypsin digested, followed by acetyl-lysine enrichment and MS analysis. Stoichiometry quantification is assessed by comparing the SILAC ratio across the dilution series. Validation of this approach was performed using AQUA peptides, which agreed with their stoichiometry calculation^30^, when stoichiometries were less than 10%. Additionally, the authors demonstrate higher quantification accuracy for peptides with very low stoichiometry (< 1%) and higher error rates with higher stoichiometry (> 1%). Therefore, the two strategies developed for quantifying acetylation stoichiometry by Choudhary *et al.* and our current study, represent orthogonal methods for accurately quantifying low and high acetylation stoichiometry, respectively.

### Acetylation dynamics and mechanistic implications

In this study, rapid acetylation dynamics were investigated using serum stimulation to initiate a major transcriptional, translational, and metabolic response^49^. Analysis in two different human cell lines revealed that acetylation changes occur on time-scales (minutes) that rival those observed via phosphorylation-dependent signaling^50, 51^. Rapid changes in acetylation included groups of sites that increased and groups of sites that decreased, many with similar kinetic profiles. Notably, many other sites showed no significant change across the four-hour time-course. The majority of acetylation sites that decreased rapidly were from proteins that increased in abundance, as would be predicted from newly translated, unacetylated proteins under serum stimulation^52^. Most importantly, lysine sites that exhibited rapid increases in acetylation are candidates for acetyltransferase control and functional regulation. The results presented here, enabled by a robust MS method, provide a critical resource for investigators studying the regulation of these pathways.

### Acetylation stoichiometry distribution in the cell

One of the most interesting observations from this study is that the highest stoichiometry and dynamics occur on proteins that exist in subcellular compartments where acetyltransferases are known to reside. In continually serum-fed MCF7 cells, we observed a significantly larger proportion of proteins with higher acetylation stoichiometry within the nuclear compartment (**Figure 4C**), which was corroborated by a separate method (**Figure 4D**). Additionally, nuclear-localized proteins displayed the most dramatic changes upon serum stimulation. Nuclear localized proteins, such as transcriptional and post-transcriptional processing factors, exhibited rapid increases in acetylation in both MCF7 and HCT116 cells. For example, we observed increases in acetylation stoichiometry ranging from 8-30% over the first hour on several splicing factors such as U4/U6.U5 tri-snRNP associated protein 1 (SART1) at K147, RNA binding-protein 8A (RBM8A) at K114, and heterogeneous nuclear ribonucleoprotein H3 (HNRNPH3) at K97. Given that these proteins display increased acetylation under pro-growth conditions and are localized to the nucleus where acetyltransferases reside, we predict that these sites are enzymatically regulated and functionally important for increasing basal splicing rates or for preferentially affecting certain types of splicing reactions. Our data provide a novel roadmap for investigating nuclear acetylation of non-histone proteins and suggest that acetyltransferases play a critical function well beyond histone acetylation.

Outside the nucleus, we observe dynamic acetylation across other cellular pathways known to be regulated during serum stimulation. We observed high enrichment for proteins involved in translation, including increases in acetylation on ribosomal proteins such as RPL3 at K103 and K155 and RPL7 at K77 (**Figures 6D and 6E**). Pro-growth increases in protein synthesis is a major cellular response to serum stimulation^52^. A known mark of actively translating ribosomes is Ribosomal protein S6 phosphorylation. The acetylation of S6 mimicked the pattern of the MCF7 cluster showing increased but slower acetylation kinetics (**Figures 5C and 5D**)^53^. Interestingly, this same cluster of acetylation sites was enriched for the Ribosome. Taken together with the contrasting observation that most newly translated proteins are unacetylated, increased acetylation of ribosomal proteins is positively associated with translation and would be predicted to be catalyzed by acetyltransferases^54^.

Another cellular pathway known to be affected by serum stimulation is mitochondrial fission and fusion, in which mitochondria transition between being highly fragmented in the quiescent state to more tubular structures as cells approach the G1-S transition^55^. On one protein involved in this process, mitochondrial fission protein 1 (FIS1) we observed large, rapid increases in acetylation at K89. FIS1 is localized to the cytoplasmic face of the mitochondrial outer membrane, therefore this site is likely to be enzymatically regulated by cytoplasmic acetyltransferases. Acetylation of FIS1 K89 might inhibit mitochondrial fission, potentially through disrupting interactions with other fission machinery such as Dynamin-1-like protein (DNM1L)^56^. Lastly, we observed dynamic protein acetylation on proteins related to protein homeostasis, particularly ER-localized proteins. It was recently shown that the lysine acetyltransferases, NAT8/NAT8B, are localized in the lumen of the endoplasmic reticulum and function to acetylate properly folded proteins as proteins traverse through the secretory pathway^57^. Acetylation by NAT8/NAT8B is proposed to signal correctly folded proteins and function in quality control. Collectively, the strong trend in these results suggest that dynamic protein acetylation occurs in subcellular compartments with known, localized acetyltransferases. The existence of bona fide protein acetyltransferases in the mitochondrial matrix is the subject of debate^9–11^. While acetylation is prevalent in mitochondria^5, 35, 36, 58, 59^, the considerably lower mean stoichiometry and no apparent kinetic trends in response to growth factor stimulation suggest that widespread enzyme-catalyze acetylation is not a major mechanism for matrix proteins. The method and results described in this study provide a valuable resource to investigate regulatory acetylation and as a rich data set for investigators to now directly test the role of acetylation in a pathway of interest.

## ASSOCIATED CONTENT

### Supporting Information

The following files are available free of charge.

Supplemental Figures – overview of the data processing pipeline, isotopic correction improves quantification of high stoichiometry acetyl-lysines, and S6 ribosomal protein S235/236 phosphorylation immunoblot (PDF).

Supplemental Table 1 – HSC Stoichiometry Curve – calculated stoichiometry and linear modeling (.xlsx)

Supplemental Table 2 – Untreated MCF7 calculated fragment ion and acetylation site stoichiometry (.xlsx)

Supplemental Table 3 – Serum Stimulated MCF7 cells calculated fragment ion and acetylation site stoichiometry (.xlsx)

Supplemental Table 4 – Serum Stimulated HCT116 cells calculated fragment ion and acetylation site stoichiometry (.xlsx)

The mass spectrometry raw files, spectral libraries, and the result files from MaxQuant and Spectronaut used in this study have been deposited to the ProteomeXchange Consortium via the MassIVE partner repository and can be accessed through either the ProteomExchange dataset identifier PXD014453 or MassIVE ID MSV000084029.

The R code used to calculate acetylation stoichiometry from the Spectronaut result files and perform all secondary analyses have been published to GitHub through Zenodo and can be accessed using the following DOI: 10.5281/zenodo.3360892.

## AUTHOR INFORMATION

### Author Contributions

The manuscript was written through contributions of all authors. All authors have given approval to the final version of the manuscript. ‡These authors contributed equally.

### Notes

The authors I.L., T.G., O.M.B., and L.R. are employees of Biognosys AG (Zurich, Switzerland). Spectronaut is a trademark of Biognosys AG.

## ACKNOWLEDGMENT

We would like to thank Greg Barrett-Wilt and Greg Sabat at the University of Wisconsin-Madison Biotechnology center for use of the Mascot database server.

## ABBREVIATIONS

ANOVA: analysis of variance
ACN: acetonitrile
DDA: data-dependent acquisition
DIA: data-independent acquisition
DDT: dithiothreitol
GO: gene ontology
HCD: higher energy collisional dissociation
HPRP: high pH reverse phase
IAA: iodoacetamide
KAT: lysine acetyltransferase
KDAC: lysine deacetylase
NCE: normalized collision energy
PSM: peptide spectrum matches
QSSA: quantitative site set functional score analysis

