## Supplemental Figures for "Revealing dynamic protein acetylation across subcellular compartments"

**Materials Included:**

1. Figure S1 – Acetylation stoichiometry data processing workflow
2. Figure S2 – Isotopic correction improves quantification of high stoichiometry acetyl-lysines
3. Figure S3 – Serum stimulation increases Ribosomal Protein S6 Ser235/236 phosphorylation
4. Table S1 – HSC Stoichiometry Curve - calculated stoichiometry and linear modeling
5. Table S2 – Untreated MCF7 calculated fragment ion and acetylation site stoichiometry
    a. Table S2_1 – MCF7 - calculated fragment ion stoichiometry
    b. Table S2_2 – MCF7 - calculated acetylation site stoichiometry
6. Table S3 – Serum Stimulated MCF7 cells calculated fragment ion and acetylation site stoichiometry
    a. Table S3_1 – Fragment ion calculated stoichiometry for serum stimulated MCF7 cells
    b. Table S3_2 – Acetylation site calculated stoichiometry for serum stimulated MCF7 cells
7. Table S4 – Serum Stimulated HCT116 cells calculated fragment ion and acetylation site stoichiometry
    a. Table S3_3 – Fragment ion calculated stoichiometry for serum stimulated HCT116 cells
    b. Table S3_4 – Acetylation site calculated stoichiometry for serum stimulated HCT116 cells

**Methods**

Immunoblot Analysis

Samples were denatured in SDS loading dye and 5 min boil. 540 ng of whole cell lysate were loaded on a Bolt 4-12% Bis-Tris Plus gel (Invitrogen), followed by transfer to PVDF. Membrane was blocked with 1% milk before being blotted with either anti-Phospho-S6 Ribosomal Protein (Ser235/236) (CST # 4856) or anti-S6 Ribosomal Protein (CST #2217) 1:1,000 and imaged. Bands were quantified using Image Studio Lite (LI-COR Biosciences).

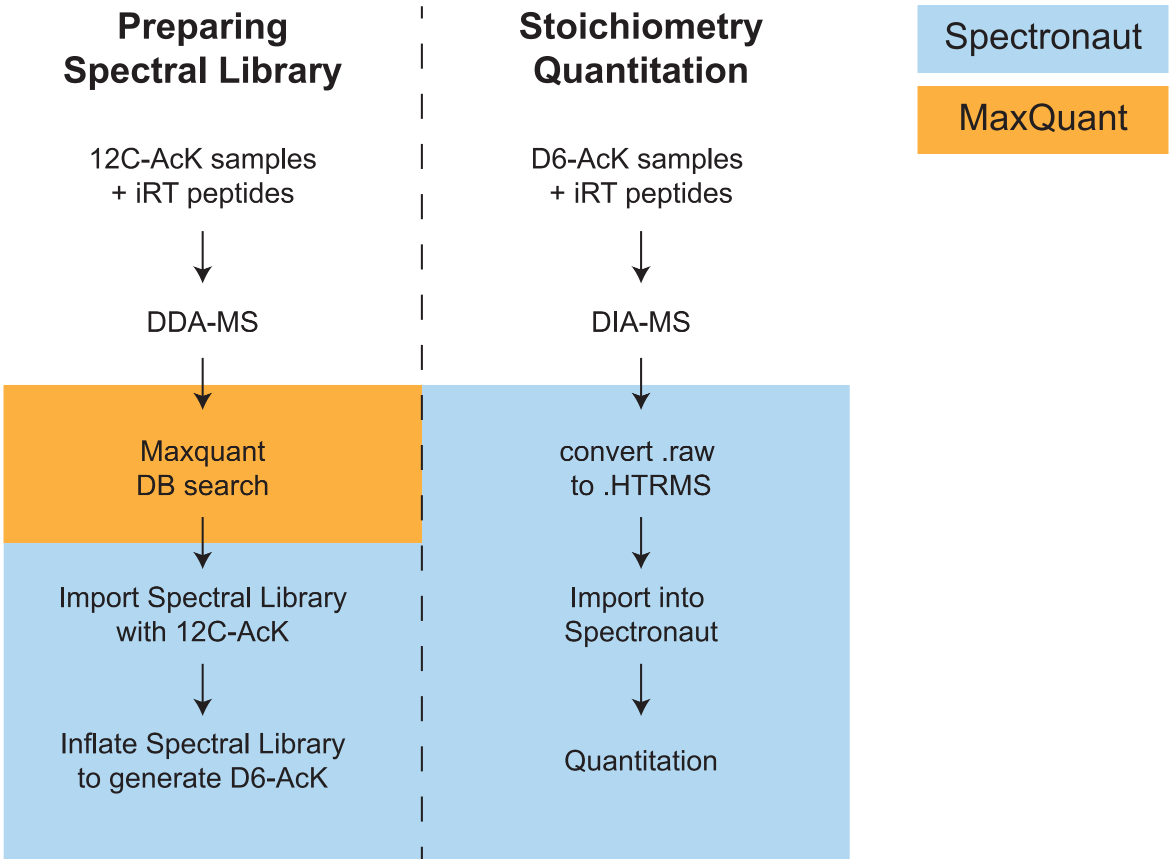


**Figure S1: Acetylation stoichiometry data processing workflow**

Diagram depicting the data analysis steps to quantify acetylation stoichiometry by generating a spectral library from DDA-MS runs in MaxQuant to deconvolute the DIA-MS runs analyzed using Spectronaut.

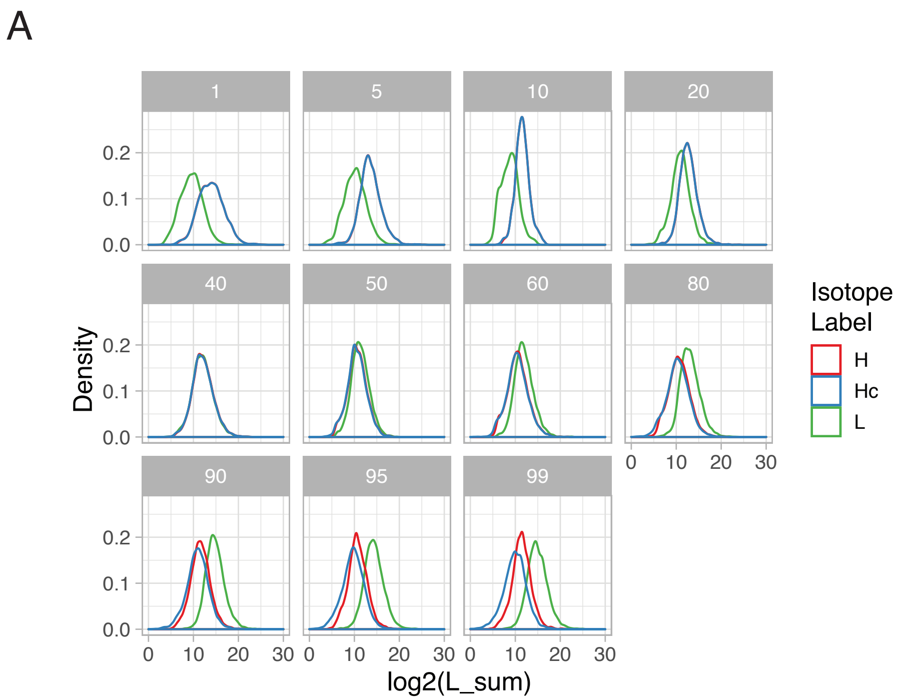


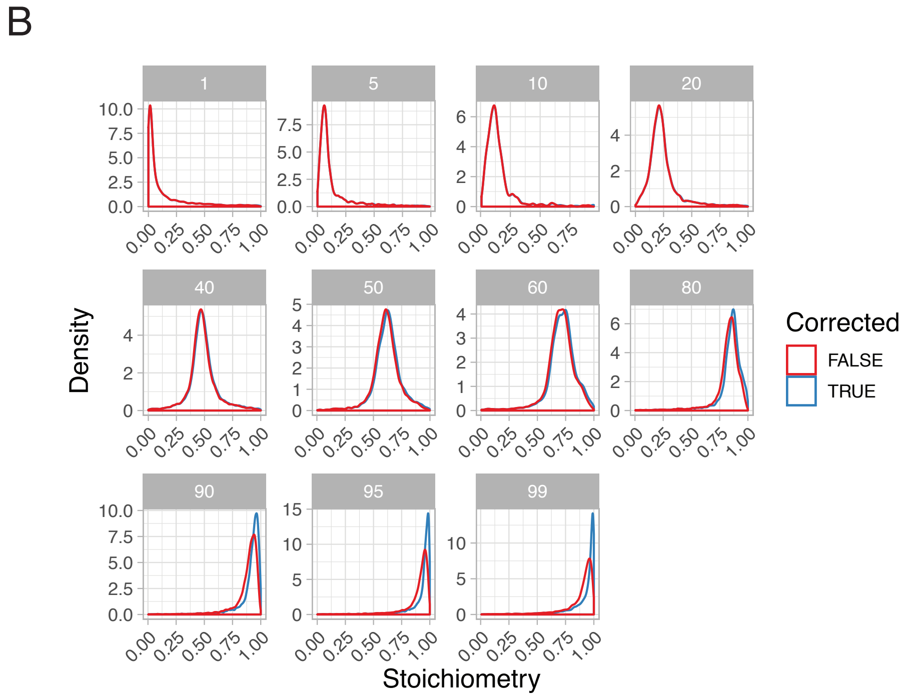


**Figure S2: Isotopic correction improves quantification of high stoichiometry acetyl-lysines**

(A) Distribution of fragment ion intensities across each input stoichiometry. Colors represent the isotopic label: green for light labeled, red for heavy labeled, and blue for heavy label that has been corrected based on the fragment ion isotopic distribution. (B) Distribution of calculated stoichiometry across each input stoichiometry. Red indicates the stoichiometry calculated from the observed fragment ion intensities and blue indicates the stoichiometry calculated from the isotopically corrected fragment ion intensities.

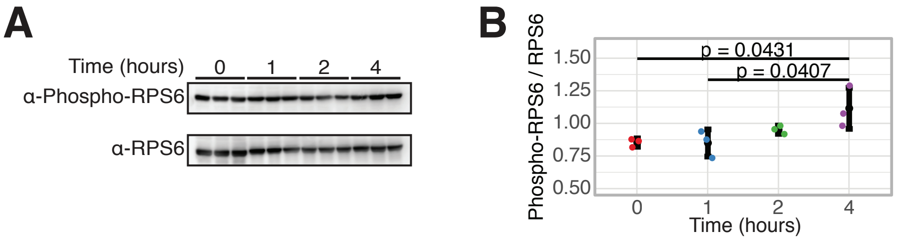


**Figure S3 – Serum stimulation increases Ribosomal Protein S6 Ser235/236 phosphorylation**

(A) Western blot of phosphorylated RPS6 Ser235/236 (top) and RPS6 (bottom). (B) Quantitation of the western blot. Statistical analysis was performed using a t-test.
